# Clinical Phenotype Prediction From Single-cell RNA-seq Data using Attention-Based Neural Networks

**DOI:** 10.1101/2023.03.31.532253

**Authors:** Yuzhen Mao, Yen-Yi Lin, Nelson K.Y. Wong, Stanislav Volik, Funda Sar, Colin Collins, Martin Ester

**Author notes:** These authors contributed equally to this work (Yuzhen Mao, Yen-Yi Lin).

## Abstract

**Motivation:** A patient’s disease phenotype can be driven and determined by specific groups of cells whose marker genes are either unknown, or can only be detected at late-stage using conventional bulk assays such as RNA-Seq technology. Recent advances in single-cell RNA sequencing (scRNA-seq) enable gene expression profiling in cell-level resolution, and therefore have the potential to identify those cells driving the disease phenotype even while the number of these cells is small. However, most existing methods rely heavily on accurate cell type detection, and the number of available annotated samples is usually too small for training deep learning predictive models.

**Results:** Here we propose the method ScRAT for clinical phenotype prediction using scRNA-seq data. To train ScRAT with a limited number of samples of different phenotypes, such as COVID and non-COVID, ScRAT first applies a mixup module to increase the number of training samples. A multi-head attention mechanism is employed to learn the most informative cells for each phenotype without relying on a given cell type annotation. Using three public COVID datasets, we show that ScRAT outperforms other phenotype prediction methods. The performance edge of ScRAT over its competitors increases as the number of training samples decreases, indicating the efficacy of our sample mixup. Critical cell types detected based on high-attention cells also support novel findings in the original papers and the recent literature. This suggests that ScRAT overcomes the challenge of missing marker genes and limited sample number with great potential revealing novel molecular mechanisms and/or therapies.

## Introduction

Accurate prediction of clinical phenotypes for patients in given cohorts is critical in advancing diagnosis, prognosis, and therapy (Ching *et al*., 2018). Heterogeneous clinical symptoms may lead to ambiguous predictions (Morley *et al*., 2021), and therefore the analyses based on high-throughput omics data have started to enter clinical routine in the last decade (Chen *et al*., 2021). A challenging step in these analyses is to dissect cellular content from patients’ genomic profiles, including the detection of cell types defined by gene expression profiles and their proportions in different patients (Newman *et al*., 2019). While clinical phenotype information, such as tumor metastasis, disease stage, and treatment response for bulk tissue samples are widely collected from various consortia, their gene expression profiles are measured by averaging cells across the whole tissue, which often do not reveal the full complexity of diverse cell types within patients.

Recent advances of single-cell and single-nuclei RNA-Sequencing (sc/snRNA-Seq) enable gene expression profiling at the unprecedented single-cell resolution. While this technology improves our understanding of cell-type markers and disease-specific signatures, analysis of large-scale cohorts is not clinically practical, especially for cancer research, for the following reasons. (1) Dependence of accurate cell type identification which might be biased or unavailable. Most scRNA-seq analysis starts with detecting cell types using unsupervised clustering, followed by cell-type annotations based on marker genes. Clinical phenotypes are then predicted by distributions of cell types or identifying specific types. However, accurate cell type identifications are affected by the marker gene information that might be suboptimal or missing, and the proper clustering resolution for a sample. Therefore, many existing scRNA-seq analysis methods even require users to provide the number of cell types, which is unknown before analysis, or set up a universal value to provide the corresponding analysis results. (2) Limited number of samples. Most well-annotated scRNA-seq datasets involve few than 20 samples, whose statistical power is too weak to support the phenotype prediction and the findings of phenotype-specific cell types. Such small size of samples can also lead to serious overfitting for most machine learning models and significantly affect their prediction performance. (3) Lack of interpretability. Many computational methods try to resolve the above issues but rarely provide users much insight into cell types and molecular mechanisms driving or related to phenotypes. Due to the above reasons, scRNA-seq often needs to be integrated with bulk assays in the analysis, and people mainly apply these methods to study compositions of single tissues rather than their clinical phenotypes for diagnosis and prognosis applications.

Here we present ScRAT, a clinical phenotype prediction framework that can learn from limited numbers of scRNA-seq samples with minimal dependence on cell-type annotations. Compared to most available scRNA-seq analysis algorithms that model gene expression profiles of different cell clusters by separate Gaussian distributions (He *et al*., 2021; Zeng *et al*., 2022), the first contribution in ScRAT is that we utilize the attention mechanism to measure interactions between cells as their correlations, or attention weights. For each cell, we incorporate all of its interaction patterns and attention weights to establish its connections with the corresponding phenotypes. Secondly, we introduce a mixup module in our framework as a data augmentation approach to mitigate the potential overfitting issue caused by the high model capacity together with the very limited number of labeled samples. Lastly, ScRAT establishes the connection between the input (cells) and the output (phenotypes) of the Transformer model simply using the attention weights. This is cost-effective compared to existing approaches from literature that tend to be computationally expensive, such as gradients propagation or training probing classifiers (Chefer *et al*., 2021b,a; Clark *et al*., 2019). ScRAT hence selects cells containing the most discriminative information to specific phenotypes, or *critical cells*, using their attention weights. It provides a natural way to construct phenotype-specific subpopulations in clinical cohorts that suggests prognostic markers and potential therapeutic information.

We evaluate ScRAT on three public COVID datasets compared to five baseline frameworks. In each dataset, we would split the samples into two clinical phenotypes based on the given annotation: COVID vs non-COVID, mild/moderate vs severe/critical, or convalescence vs progression. We also reduce the number of training samples to investigate the predictive power of each framework. ScRAT achieves the best AUC in all comparisons and provides leading precision and recall in most scenarios. The performance edge of ScRAT over its competitors increases as the number of training samples decreases, indicating the efficacy of our sample mixup module. Since these public datasets come with cell type annotations in various resolutions, we also examine the connections between phenotypes and subpopulations enriched with high-attention cells. What’s more, our experiment shows that ScRAT can detect disease-critical and phenotypic-driver subpopulations using high-attention cells, and these information can potentially help to identify novel conditions of druggable populations.

In short, ScRAT is the first deep neural network based method to predict clinical phenotypes from scRNA-Seq, and among the first attention-based framework for scRNA-seq analysis. Our integration of attention mechanism and mixup allows ScRAT to be independent of cell-type annotations, capable of learning from a limited number of training samples. Lastly, we propose a simple method to explain the prediction of Transformer that is more cost-effective than the existing methods. This indicates ScRAT can provide interpretable information to guide biologists.

## Related Work

### Deep Learning in Single-cell RNA-seq Analysis

Single-cell RNA-seq has become a popular tool for gene expression analysis at a single-cell resolution. However, analyzing scRNA-seq data is a challenging task, and traditional bioinformatics methods may not be able to handle the complexity and heterogeneity of the data. Deep learning techniques have been applied to scRNA-seq analysis, showing promising results on many related tasks. For example, Yin *et al*. (2022) propose an autoencoder-based classification framework to obtain compressed representations of scRNA-seq data. These representations are then fed into subsequent classifiers to predict the cell types. Ravindra *et al*. (2020) use graph attention networks (GAT) to construct a graph representation of the scRNA-seq data, where each node represents a cell and each edge represents the similarity between two cells. Then the disease state for each cell is predicted based on the learned graph representations.

### Phenotype Prediction Using Bulk-cell RNA-Seq

Gene expression profiling has been used to predicting phenotypes in many clinical settings (Lonsdale *et al*., 2013; Uhlen *et al*., 2017). PAM50 classifies breast tumor based on expression profiles of 50 genes (Perou *et al*., 2000). Molecular phenotypes of prostate cancer also relie on gene expression profiling (Cancer Genome Atlas Research Network, 2015), and multiple expression-based diagnosis tests have also been developed. For example, The Prolaris cell-cycle progression (CCP) predicts aggressiveness for prostate cancer using expressions of 31 genes from the cell cycle proliferation pathway (Cuzick *et al*., 2012). A signature of 157 genes was developed to predict lethal prostate cancer (Penney *et al*., 2011). Oncotype Dx genomic prostate score (Cullen *et al*., 2015) and Decipher Biopsy score (Erho *et al*., 2013) also identify gene signatures to predict the risk of metastasis as the tumor outcome. These methods are mainly developed using bulk assays, and can not benefit from cell-level resolution information in scRNA-seq to improve diagnosis and prognosis.

### Phenotype Prediction Using Single-cell RNA-Seq

CloudPred (He *et al*., 2021) models the individual points as samples from a mixture of Gaussians, probabilistically assigns points to clusters, then estimates prevalence of the subpopulations and use it to predict the phenotype of that patient. scPheno (Zeng *et al*., 2022) constructs gene expression profiles by a joint distribution of cell states and disease phenotypes based on a deep generative probabilistic model, and feeds the distribution as the predictive features to support vector machine (SVM) for the phenotype prediction. One of the main weaknesses of these methods is that neither of them uses deep neural networks, which indicates limited model capacity, although it also allows these methods to work under a limited number of labeled training data. Besides, the assumption of modeling single cell population as Gaussian is doubted, and useful information might be ignored in that case.

## Problem definition

A *cell* is the most basic unit in scRNA-seq experiments and will be denoted by a vector c over m genes including the measure of gene expression level, e.g. the count of Unique Molecular Identifiers (UMIs). The ultimate prediction unit is denoted as a *sample*, which is extracted from a single patient. A sample consists of n cells and is represented as an *n* × *m* matrix *S*, where *S*_*ij*_ corresponds to the raw or normalized UMIs of the *j*-th gene in the *i*-th cell. Each sample is associated with a specific one-hot encoded phenotype label from a pre-defined set *P* = {*P*^1^, *P*^2^, ··· *P*^*o*^}. Based on that, we formally define our problem as follows:

### Problem

Phenotype prediction for scRNA-seq samples.

### Given

A set of labeled samples represented by scRNA-seq matrices *D* = {*S*_1_, *S*_2_, ···, *S*_*K*_}, and their corresponding labels *Y* = {*P*_1_, *P*_2_, ···, *P*_*K*_}; a set of unlabeled samples 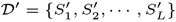.

### Find

A prediction model which maps 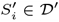 to *P*^*i*^ ∈ 𝒫.

## Method

In this section, we present a neural-network-based method called ScRAT to predict the phenotype of an scRNA-seq sample. An overview of ScRAT is presented in Fig. 1, which consists of three major modules: Sample Mixup, Attention Layer and Phenotype Classifier. Our method takes an scRNA-seq sample from a single patient as input. Note that the order of cells within each sample does not matter and the size of each sample is variable. To alleviate the possible over-fitting issue, we employ a data augmentation technique called “sample mixup” during the training time to increase the amount and diversity of training samples. The backbone of ScRAT is a multi-head attention layer (Vaswani *et al*., 2017) which aims to learn a task-orientated embedding for each cell within the sample. Considering its poor scalability (Tay *et al*., 2020), a cropping strategy is applied to the input sample before passing it to the attention layer. As the last step, a one-layer Multi-Layer Perceptron (MLP) takes the output of the attention layer and predicts the phenotype as a probability distribution over the different values of the phenotype. In the following subsections, we delve into these three modules of ScRAT in detail.

**Fig. 1.**
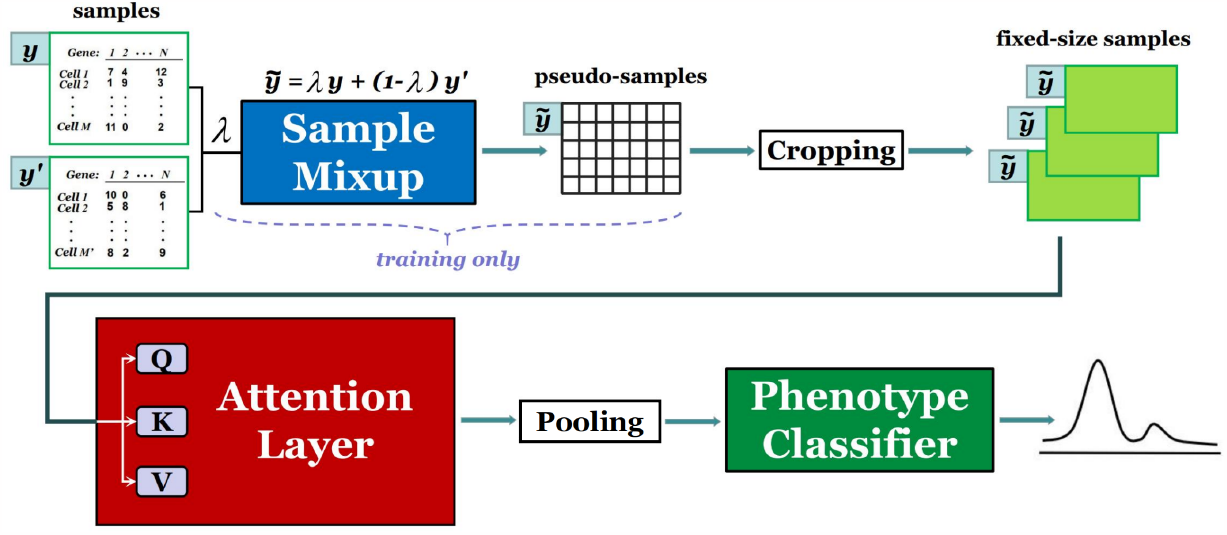
An overview of **ScRAT**, which consists of three main modules: Sample Mixup, Attention Layer and Phenotype Classifier. It takes a scRNA-seq sample (a set of cells) as input, and outputs the predicted phenotype for the input sample.

### Sample Mixup

The size of currently available scRNA-seq datasets is very small, and it is expected to remain relatively small in the near future, which will likely result in overfitting when training a deep learning model. Mixup and its variants (Zhang *et al*., 2017; Verma *et al*., 2019) are interpolation-based and widely-adopted data augmentation techniques for regularizing neural networks and improving model generalizability (Carratino *et al*., 2020). For instance, in computer vision setting, mixup convexly combines random pairs of images and their associated labels to generate new training data. Inspired by this, for scRNA-seq analysis, we introduce a simple but efficient data augmentation method, sample mixup, to generate new samples during training process. Specifically, given two scRNA-seq samples *S* and *S*^’^ together with a fixed λ ∈ [0, 1], sample mixup is defined as follows (Zhang *et al*., 2017):

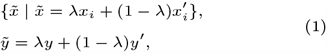

where *x*_*i*_ and 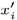 are gene expression profile of cells drawn from *S* and *S*^’^, and *y* and *y*^’^ are corresponding one-hot phenotype label encodings.

Compared to the computer vision setup, samples here correspond to images, cells in each sample correspond to pixels in each image, and phenotypes of samples correspond to labels of images. The main differences between these two scenarios are that pixels in one image can only be mixed with pixels in the same spatial location of another image, and mixup can only be applied to images having the same size. scRNA-seq data is not limited by these two constraints.

The proposed scRNA-seq sample mixup aims to increase the number and diversity of samples. Specifically, given a pair of samples *S*_1_ and *S*_2_ with the same or different phenotypes, we first randomly sample a batch *S*_11_ containing *N* cells only from *S*_1_, and sample another batch *S*_21_ with the same amount of cells only from *S*_2_. Each batch is allowed to include duplicate cells during sampling. Then mixup is applied to *S*_11_ and *S*_21_ based on Eq. 1, where λ ∼ Beta(*α, α*), for *α* ∈ (0, ∞) (Zhang *et al*., 2017; Carratino *et al*., 2020), to generate *N* augmented cells forming a new sample *S*_3_ called pseudo-sample, with the phenotype label equals to the linear combination of phenotype labels of *S*_1_ and *S*_2_.

Notably, since cells of different populations are biologically very different, it does not make much sense to directly apply mixup to them. Therefore, although our model does not require cell type information, during the sample mixup, we only mix cells of the same cell population, assuming that this information has either been annotated by a human expert or been determined automatically by state-of-the-art annotation methods such as MARS (Brbić *et al*., 2020). For cell populations that appear only in one of the samples, we add Gaussian noise to the gene expression profile of cells that belong to those unique cell populations during the mixup.

Sample mixup also ensures that the proportion of each cell population in the pseudo-sample is the linear combination of the proportions of that cell population in two original samples. For example, given λ = 0.2 and the proportions of cell population A in two original samples are 30% and 20% respectively, then the proportion of A in the pseudo-sample is calculated as: 0.2 × 30%+0.8 × 20% = 22%.

The effectiveness of sample mixup has been evaluated in our ablation study (Section 5.2.1).

### Attention Layer

Attention mechanisms (Bahdanau *et al*., 2014) have achieved state-of-the-art performance in a wide range of machine learning tasks which take a set of elements as input, such as words (Devlin *et al*., 2018) and pixels (Dosovitskiy *et al*., 2020). An attention mechanism pays more attention to the relatively important elements by assigning high weights to them during the forward pass. Multi-head Attention is one of the most popular version of this mechanism which was first proposed in (Vaswani *et al*., 2017), and we use attention as a synonym for this version here. Compared with classical neural nets such as MLP and CNN (Krizhevsky *et al*., 2017), attention can not only deal with variable-sized inputs but also assign weights to different elements dynamically, which is necessary for unordered inputs.

Specifically, the input of the attention layer is a set of cell embeddings 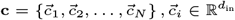, where *N* is the number of cells, and *d*_in_ is the number of features in each embedding. Following the previous work (Vaswani *et al*., 2017) closely, our attention layer maps the input embeddings to three different kinds of vectors: key, query and value using three weight matrices with the same shape respectively: 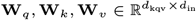, where *d*_kqv_ is the dimensionality of key, query and value. Afterwards, a self-attention with scaled dot-product is applied to each pair of cells to compute their attention weights based on their key and query vectors:

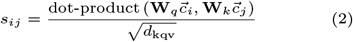

which denote the importance of cell *j* to cell *i* and are normalized using the softmax function:

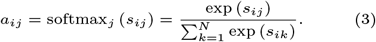

These attention weights are then treated as the weights in the following linear combination process which outputs a new embedding for each cell based on the value vectors of all cells:

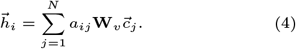

To extract information at different positions as well as make the training process more stable (Liu *et al*., 2021), multi-head attention (Vaswani *et al*., 2017) is applied in our attention layer. Specifically, instead of utilizing only one attention head with one group of **W**_*q*_, **W**_*k*_, **W**_*v*_, we utilize *K* attention heads with *K* different groups of mapping matrices and run them in parallel. Afterwards, we concatenate the outputs from each head and apply an additional linear layer to it at the end.

In a nutshell, our attention layer is formulated as:

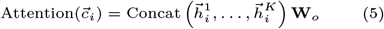

where 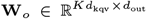 is a weight matrix, 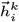 is the output of the *k*-th head based on Eq. 4.

One limitation of existing attention-based models is that they cannot handle very long sequences as input since the self-attention operation has quadratic run-time and memory complexity (Beltagy *et al*., 2020; Zhou *et al*., 2021). Therefore, after augmenting the whole dataset using mixup, we introduce a cropping strategy to both training and test data, which randomly selects several subsets from each sample and only use these subsets to train the model. We call these subsets of cells “fixed-size samples” in this paper. More specifically, for each sample, we randomly select *NC* cells as one fixed-size sample, and generate *NS* fixed-size samples for each sample. During the training process, each fixed-size sample is calculated a loss which is added up in the final loss computation used to update the model parameters; while during the testing process, we assign a (categorical) predicted label to each fixed-size sample by setting a threshold, and use majority vote to assign the predicted label to each sample based on their fixed-size samples. Here, *NC* and *NS* are both hyper-parameters which can be tuned by the users. Since *NC* could be relatively small, this cropping strategy improves the model scalability. Moreover, this strategy is an analogy to the cropping in the computer vision setting, and therefore can also be treated as a useful data augmentation approach. In the next section, comprehensive experiments demonstrate its effectiveness.

### Phenotype Classifier

The output of the attention layer are the embeddings of all cells within the input sample. Similar to the way the average pooling function operates in image classification, we aggregate the cell embeddings for each sample by computing the average value along each dimension. While this method may cause some loss of information, it is a commonly used and effective technique to simplify the feature map representation and improve the model’s generalization performance. Moreover, it ensures that the cell order does not affect the final results. Finally, the aggregated embedding is passed to the phenotype classifier, a one-layer MLP, which outputs the predicted phenotype for the input sample, i.e. a probability distribution over the different values of the phenotype.

## Experiments

We evaluate the performance of ScRAT on three large-scale public COVID scRNA-seq datasets, and compare it with five state-of-the-art methods. We perform an ablation study to determine the impact of the different ScRAT components. Finally, we design a cost-effective method to convert cell attention weights in ScRAT as the relevance score which determines the relevance of a given cell population with respect to the clinical phenotype. Our biological analysis demonstrates the potential of revealing disease mechanisms based on the critical cell types identified using attention weights in ScRAT.

### Experimental Setup

#### Datasets

Our experiments include four tasks based on the following three scRNA-seq COVID19 cell datasets. For COMBAT (COvid-19 Multi-omics Blood ATlas (COMBAT) Consortium, 2022) and Haniffa (Stephenson *et al*., 2021) datasets, we perform the task of disease diagnosis (i.e., COVID vs Non-COVID). For SC4 (Ren *et al*., 2021) which includes mostly COVID samples, we perform two separate tasks of predicting severity (i.e., mild/moderate vs severe/critical) and stage (i.e., convalescence vs progression). See Table 1 for more information.

**Table 1.** Summary of 3 Datasets. We exclude any samples with less than 500 cells or unclear clinical phenotype annotations from the original datasets. Note that one patient can be sampled multiple times and contribute to multiple samples. For Combat and Hannifa, the numbers of samples in Class 1 and 2 correspond to the number of COVID and non-COVID samples. Class 1 and 2 in SC4 correspond to mild/moderate vs severe/critical phenotypes among 91 samples of progression for the severity prediction, and convalescence vs progression for the stage prediction. See Supplementary Table 1 - 3 for detailed information.

| Dataset | Phenotype | #Cells | #Samples | #Samples in Class 1 | #Samples in Class 2 |
| --- | --- | --- | --- | --- | --- |
| Combat | Disease Diagnosis | 835,937 | 121 | 77 | 44 |
| Haniffa | Disease Diagnosis | 528,438 | 105 | 71 | 34 |
| SC4 | Disease Severity | 501,943 | 91 | 29 | 62 |
|  | Disease Stage | 1,289,496 | 229 | 138 | 91 |

#### Design of Experiments

To reflect the limited number of labeled scRNA-seq samples in real applications, we first define the *Training Ratio* for each task as the number of samples included in the training data divided by total number of samples in the dataset, and split the original dataset into the training and test datasets accordingly. For each given training ratio ranging from 9% to 50% in our design, we run the experiments for 100 random splits to better evaluate the performance of different methods. Considering the high dimensionality of scRNA-seq data which is likely to result in serious over-fitting, we map the original input to a low dimensional latent space and keep only 50 principal components by PCA (Halko et al., 2011). The area under the receiver of characteristic curve (AUC) is used as the evaluation metric in the following discussions.

#### Baselines

We compare ScRAT with five popular phenotype prediction methods, including two pseudo-bulk methods and three single-cell methods. For pseudo-bulk methods, we first average the gene expression across all cells in one sample to simulate a pseudo bulk assay as the input to the prediction model. We choose naive linear layer and feed-forward layer as two such prediction models which are denoted as **Linear** and **Feedforward (bulk)** respectively in this paper. For single-cell methods, single-cell resolution information can be used by two different strategies, either processing each cell separately, or processing all cells in one sample interactively. Simple models such as linear layer and feed-forward layer can be only used for the first strategy since their weights are position-specific, the prediction results change according to the order of cells, which makes it difficult to process multiple cells as a whole. In the experiment, we use feed-forward layer for this strategy and denote this baseline as **Feedforward (single)**. For the second strategy of interactively analyzing all cells, the encoding of a given cell will be affected by others in the same sample and can potentially capture correlations between cells. Vanilla attention layer (Vaswani *et al*., 2017) and CloudPred (He *et al*., 2021) are selected as the methods for this strategy, which are denoted as **Attention** and **CloudPred**. We set 10 as the number of clusters in Cloudpred since it achieves the highest AUC among most of the experiments compared with other numbers of clusters we try (5 and 20). Notably, baseline Attention is equivalent to using the ScRAT without sample mixup module.

#### Configuration of ScRAT

Throughout our experiments, we observed that the performance of the model remained largely unaffected when the number of pseudo-samples exceeded 250 (Fig. 2). Therefore, for each experiment of ScRAT, we apply mixup to generate 300 pseudo-samples with 10,000 cells in each from the original training samples, and only use these 300 pseudo-samples to train the model. α in the beta-distribution of mixup is set to 0.5. For the cropping strategy, we set the number of cells in each fixed-size sample (*NC*) to 500, and set the number of the fixed-size samples (*NS*) to 20 and 50 for training and testing respectively. We only use one attention layer, and set the number of attention heads K to 8 and the dimension of each head d_*kqv*_ to 16. We use Adam optimizer with learning rate as 1e-2. All the hyper-parameters are decided using the 5-fold cross validation technique.

**Fig. 2.**
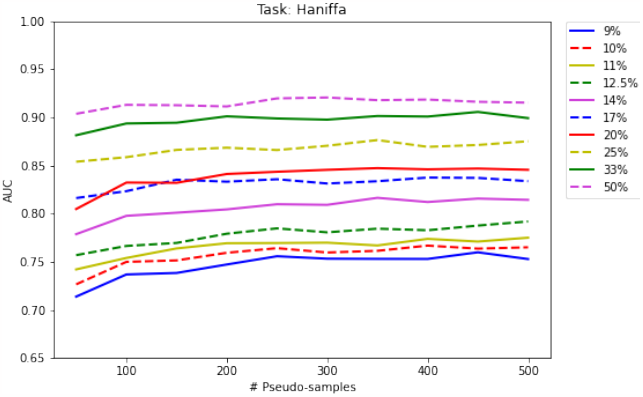
Impact of the number of the pseudo-samples on ScRAT AUC. Our experimental results indicate that ScRAT’s performance (AUC) is not significantly impacted by the number of pseudo-samples generated by Sample Mixup once it exceeds 250.

### Prediction Results

We compare ScRAT with five baseline methods on four tasks, and provide the AUC of all methods in Fig. 3. In general, we have the following observations: (1) ScRAT consistently outperforms all baseline methods on four tasks, which demonstrates the effectiveness and generalizability of ScRAT. More specifically, the performance edge of ScRAT over the second best method (usually the vanilla attention) increases as the training dataset size (number of samples) decreases, verifying the usefulness of our proposed sample mixup as a data augmentation approach. For example, at training ratio = 9%, the p-value of the t-test between AUCs of ScRAT and vanilla attention is much smaller than 0.01 in all but the SC4-Severity tasks. (2) Vanilla attention layer is the second best model for all four tasks, which indicates the strengths of the attention mechanism in phenotype prediction task using scRNA-seq data. (3) Feed-forward (single) has the highest recall in both COMBAT and Haniffa but low precision compared to the attention model and ScRAT, suggesting the necessity of simultaneously considering information from all cells (Fig.1&2 in the supplementary material).

**Fig. 3.**
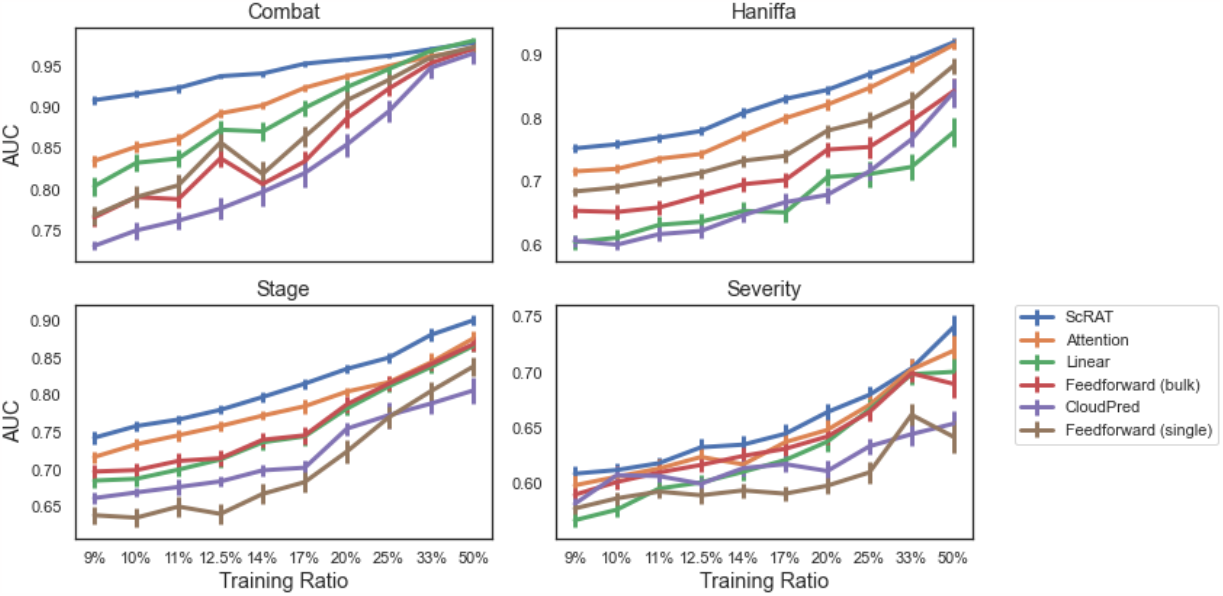
Comparison of different methods on four different tasks. For each task, we report the prediction results of all methods using AUC *±* 95% confidence intervals for 10 different training ratios. ScRAT outperforms other methods in all settings, followed by vanilla attention (The p-value of t-test between ScRAT and vanilla attention *<<* 0.01 in all but the SC4-Severity tasks at Training Ratio=9%). The performance edge of ScRAT over vanilla attention increases as the training ratio decreases, especially for the Combat datasets. See Supplementary Table 4 - 7 for detailed information.

#### Ablation Study

##### Impact of Mixup Strategies

To investigate the impacts of applying mixup only to cells from the same population, we use the Haniffa dataset to compare the performance between ScRAT (i.e., mixup of cells from the same population) and an alternative method that applies mixup to random pairs of cells regardless of their cell populations. The results are shown in Fig. 3(a). Following the predefined clusterings of 9/18/12 cell populations in Combat/Haniffa/SC4, sample mixup can improve the phenotype prediction performance and increases the AUC by up to 1.4% compared to the alternative method, which indicates the efficacy of applying mixup to cells from the same population.

What’s more, given the number of cells in these datasets, these clusterings are considered low-resolution and can be achieved automatically without human intervention. This indicates that the sample mixup of ScRAT has a minimal dependency of accurate cell type annotations, and can avoid the common challenges of finding the best resolution in scRNA-seq analysis. Moreover, our experiment shows that even without any predefined population information, the random mixup can still be applied to overcome the bottleneck of a limited number of training samples without dramatically hurting the performance.

##### Impact of Attention Weights

Despite the wide discussions (Serrano and Smith, 2019; Jain and Wallace, 2019; Wiegreffe and Pinter, 2019), the usefulness of the attention mechanism in interpretability is still controversial. Inspired by a recent work about top *k* attention weights (Gupta *et al*., 2021), we provide a new perspective for the attention mechanism interpretation by building a bridge between attention weights and the model performance. We design the following two experiments to empirically achieve this goal: given a sample with *N* cells, we modify the *N* × *N* self-attention matrix described in Eq. 3 in two different ways:

<u>Top</u> *<u>k</u>*: For each row of the matrix, we only keep entries with top *k* largest attention weights, denoted as (*a*_1_,…, *a*_*k*_), and set attention values of all remaining entries uniformly as 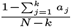.

<u>Random</u> *<u>k</u>*: For each row of the matrix, we only keep attention weights for *k* randomly selected entries, denoted as 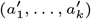, and set the attention values of remaining entries uniformly as 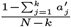.

The results of the above two experiments on Haniffa dataset are presented in Fig. 4(b), where we set *k* to 5. The AUC of the top *k* attention method is almost the same as the vanilla attention, which indicates that the top *k* attention weights are sufficient for the prediction task. On the other hand, the performance of the random k method drops significantly, suggesting the necessity of keeping high attention weights. In this way, we empirically prove the connections between the attention weights and the model performance, and provide an attention mechanism interpretation for ScRAT.

**Fig. 4.**
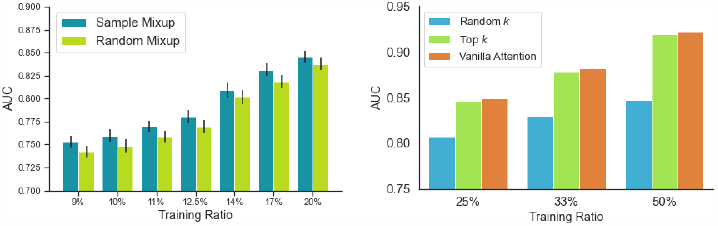
Ablation Study. **Left(a):** *Sample Mixup* corresponds to the current procedure of ScRAT that only mixes cells from the same population. *Random Mixup* means that we apply mixup to any pairs of cells. The result indicates the validity of current sample mixup method. **Right(b):** We compare the performance of various attention matrix re-construction strategies to show that attention weights are related to the model performance. For each row of the attention matrix, *Top k* keeps the top *k* largest attention weights, and *Random k* keeps *k* randomly selected attention weights, before normalizing the matrix for the remaining fields. Here, *k* is set to 5. The results of ScRAT using the original attention matrix described in Eq. 3 is denoted by *Vanilla Attention*. The result shows that top *k* method achieves AUC comparable to using all attention weights, and they are both better than random *k* attention. This provides strong evidence of the connection between high attention weights and model performance.

### Biological Interpretation of Cell Attention

The accuracy of predicting phenotypes using ScRAT relies heavily on high-attention cells, indicating a strong connection between these cells and the critical-cell-types-driving clinical phenotypes. While ScRAT calculates attention weights without cell type annotation, we use the manually annotated cell types provided by the authors of the COVID datasets to examine the biological meaningfulness of high-attention cells. We demonstrate that the most relevant cell types among the high-attention cells, such as specific monocytes and platelets, support findings in the original paper and the recent literature. Our analyses confirm the relevance of high-attention cells to clinical phenotypes, and their potential for determining critical cell types for other diseases.

We first define the relevance of a cell with respect to the phenotype prediction as follows. Given a trained model with H attention heads and an input sample *S*_*j*_ with *N* cells, ScRAT generates one attention matrix per attention head. For a cell *c*_*i*_ in *S*_*j*_, its *High-attention Occurrence Value* (HOV) is defined as the total number of times its attention weight ranked top *k* in a row across all rows and all *H* attention matrices, or

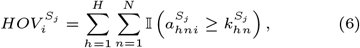

where 𝕀 (·) is the indicator function, 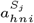 is the attention weight at the *n*-th row and *i*-th column of the *h*-th head’s attention matrix, and 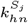 denotes the *k*-th highest attention weight in the same row of the same attention matrix.

Once we have cell annotations for all cells in *S*_*j*_, we extend the cell-level HOV to derive the *Relevance score* (R-score) for any given cell type 𝒯 with respect to sample *S*_*j*_ by adding together the HOVs of all the cells in 𝒯 ∩ *S*_*j*_, and normalize it:

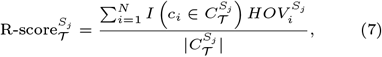

where 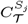 is the set of all the cells of cell type 𝒯 in *S*_*j*_.

For every phenotype, we then average the R-scores of the same cell type across all samples of this phenotype. The top *k*^’^ cell types with the highest averaged R-scores are then selected as the critical cell types of this phenotype.

Here we use the Haniffa dataset, the most comprehensively analyzed one among the three datasets used in the experiments, to demonstrate the clinical relevance of high-attention cells and critical cell types reported by ScRAT. The top 10 critical cell types (among 51 cell groups defined in the original paper) ranked by R-score are shown in Table 2. Assuming critical cell types would better separate patients of different phenotypes, we test how well a phenotype can be predicted using only cells from that cell type and a simple feed-forward network. The AUC reported in Table 2 corresponds to a 50% training ratio. Most of our critical cell types have AUC > 0.85. We repeat the experiment for all 51 cell types and the AUC of critical cell types selected by ScRAT are among top 10 AUC except for RBC and pDC, which demonstrates the relevance of cell types with a high R-score for phenotype prediction.

**Table 2.** Top critical cell types with their R-score and AUC of phenotype classification when using only the corresponding critical cell type. We rank these critical cell types based on their R-score of COVID phenotype. The larger R-score indicates the higher relevance to the phenotype. We use the Feedforward (single) model to predict the phenotype using only cells from single cell type. The AUC is based on 50% training ratio, using half of patients as the training data and the other half of patients for testing. Most critical cell types selected by ScRAT also achieve high AUC except for RBC and pDC. A cell type of higher AUC is more discriminative in predicting different phenotypes, and hence more likely a real critical call type. The concordance between cell types with high R-scores and high AUCs shows that high-attention cells detected by ScRAT are phenotype-specific. See Supplementary Table 8 for detailed information.

| Critical Cell Types | R-score | AUC |
| --- | --- | --- |
| pDC | 2.58 | 0.58 |
| RBC | 2.20 | 0.68 |
| Plasmablast | 2.13 | 0.94 |
| Platelets | 2.13 | 0.86 |
| HSC_CD38pos | 1.69 | 1.00 |
| Plasma_cell_IgG | 1.62 | 0.95 |
| CD83_CD14_mono | 1.56 | 0.89 |
| CD14_mono | 1.50 | 0.94 |
| Plasma_cell_IgA | 1.24 | 0.92 |
| DC3 | 1.18 | 0.77 |

Next we compare these critical cell types to the corresponding analysis in the original paper (Stephenson *et al*., 2021), and discover that their major findings are related to the critical cell types listed in Table 2: (1) **Humoral immune response**. ScRAT detected multiple subtypes of Plasma cells as critical, i.e., Plasmablasts, Plasma_cell_IgG, and Plasma_cell_IgA, which are the key effectors of the humoral immunity that produce antibodies. Consistent with our finding, the authors of the original paper also reported a larger population of Plasmablasts, Plasma_cell_IgA, and Plasma_cell_IgG in COVID-19 patients with severe symptoms. Notably, one characteristic of the humoral response against SARS-CoV-2 is the short-lived neutralizing antibodies, both IgG and IgA, manifested in the different humoral responses during COVID-19 infection and of other inflammatory conditions (Nguyen *et al*., 2022). More than 21% of cells from these 3 subtypes have high-attention weight, suggesting that ScRAT can detect the significance of humoral immune response during COVID-19 infection. (2) **Impacts of monocytes**. ScRAT also identified CD14_mono as critical cell types. The data in (Stephenson *et al*., 2021) implies that CD14+ monocytes preferentially replenish the bronchoalveolar macrophages in health, while a much smaller and specific subset of monocytes, namely the C1QA/B/C+/CD16+ monocytes, replenish the bronchoalveolar macrophages of the COVID-19 patients. The latter denoted as C1_C16_mono ranks 13^*th*^ based on the R-score, and its population expansion is also more often observed in patients admitted to ICUs. The differential behaviors of these monocytes constitute a distinguishing feature between COVID-19 and non-COVID-19 patients. (3) **Monocytes and platelet aggregates**. Pathological monocyte-platelet interactions have been associated with aberrant coagulation and thrombosis formation in COVID-19 patients (Levi *et al*., 2020; Hottz *et al*., 2020). Since such interaction requires receptor-ligand interactions, the original authors suggested several receptor-ligand pairs between monocytes and platelets that may contribute to the aberrant interactions in COVID patients. This finding supports the selection of more than 40% of platelet cells as critical by ScRAT. (4) **Hematopoietic stem cells**. HSC_CD38pos are early hematopoietic progenitors and are rarely observed in PBMC samples. The authors hypothesized that their presence in the PBMC samples of COVID-19 patients reflected perturbations of the bone marrow homeostasis during COVID-19 infection. Since HSC_CD38pos only constitutes less than 0.27% of cells in the dataset, this demonstrates that ScRAT can detect important phenotype-specific cell types of very small size.

We also want to highlight the detection of DC3 at the bottom of Table 2. This newly identified dendritic cell type has been shown to promote inflammatory functions of CD4+ and CD8+ T cells (Villar and Segura, 2020), but their specific functions are yet to be deciphered. There are recent reports about their association with COVID, including an increased of CD163+ CD14+ cells within the DC3 cell type in COVID patients with severe symptoms (Winheim *et al*., 2021). Although the exact roles of these DC3 cells in COVID infection are yet to be uncovered, their high R-scores show the ability of ScRAT to detect cells of interest in a specific biological context.

These analyses suggest that the critical cell types reported by ScRAT are indeed phenotype-specific, consistent to, and supported by verified biological knowledge and the latest findings.

## Conclusion

In this paper, we introduce the problem of phenotype prediction using scRNA-seq data. We present ScRAT, an attention-based method that is designed to learn from limited samples without prior knowledge of marker genes or critical cell types, and provides accurate phenotype predictions. ScRAT consists of three sub-modules: Sample Mixup, Attention Layer and Phenotype Classifier. Sample Mixup increases the size of training data to avoid overfitting. The Attention Layer models interactions between cells without any given cell-type annotations, and provides a way to extract critical cells important in phenotype predictions. The Phenotype Classifier takes the latent representation of the input data produced by the attention layer and predicts the phenotype. We perform experiments on four tasks from three benchmarks and demonstrate that ScRAT consistently outperforms five baselines. We also show the biological meaningfulness of the cell types which ScRAT determines to be critical for phenotype prediction, through an analysis of the papers of the consortia which create the benchmarks and of several more recent studies. These findings suggest that ScRAT has the potential to discover phenotypic-driver cell types that suggest novel molecular mechanisms and/or targeted therapies.

## Future Work

While we use COVID datasets as our testbed due to the greater availability of public domain datasets, our ultimate goal is to predict clinical phenotypes and discover phenotype-specific cell types for cancers that are one of the most heterogeneous ecosystems with high variance among patients. In our future work, we plan to collect scRNA-seq cancer datasets to investigate the efficacy of ScRAT for diagnosis and prognosis for various types of cancer. The current design of ScRAT does not offer an easy integration of data from different cohorts, such as learning a model on data from one consortium and applying it to predict clinical phenotypes on the data of other consortium data. The challenges are differences in sequencing protocols, patient-level variance, and batch effects observed in all atlas initiatives (Luecken *et al*., 2022). Differences in analysis protocols, including cell-type annotation ontologies, create further challenges. We will explore ways to employ recent advances in transfer learning to increase the transferability of ScRAT models.

## Supplementary Meterials to ScRAT

### 1 Summary of COVID scRNA-seq Atlases

This section provides the breakdown of samples from each atlas in our experiment. A single patient can be sampled multiple times and contribute to multiple samples. Since we exclude samples with less than 500 cells, we define *effective samples* as those samples with at least 500 cells, and *effective cells* as the collections of all cells from effective samples used in our experiments.

**Table 1:** Overview of Combat scRNA-seq dataset.

| Labels | COVID status | Annotations | Number of Samples | Number of Cells | Number of Samples < 500 cells | Number of effective samples | Number of effective cells |
| --- | --- | --- | --- | --- | --- | --- | --- |
| HV | No | Healthy | 10 | 92,205 | 0 | 10 | 92,205 |
| FLU | No | Influenza Acute | 12 | 19,233 | 1 | 11 | 19,058 |
| Sepsis | No | Sepsis acute(IP) | 23 | 164,128 | 0 | 23 | 164,128 |
| COVID_MILD | Yes | Mild | 17 | 114,418 | 0 | 17 | 114,418 |
| COVID_SRV | Yes | Severe | 30 | 247,799 | 0 | 30 | 247,799 |
| COVID_CRT | Yes | Critical | 17 | 93,982 | 1 | 16 | 93,969 |
| COVID_LDN | Yes | Low-dose Naltrexone | 2 | 15,485 | 0 | 2 | 15,485 |
| COVID_HCW_MILD | Yes | Community COVID-19 | 13 | 88,898 | 1 | 12 | 88,875 |

**Table 2:** Overview of Hannifa scRNA-seq dataset.

| Labels | COVID status | Annotations | Number of Samples | Number of Cells | Number of Samples < 500 cells | Number of effective samples | Number of effective cells |
| --- | --- | --- | --- | --- | --- | --- | --- |
| Healthy | No | Healthy | 24 | 97,039 | 0 | 24 | 97,039 |
| LPS | No | LPS | 12 | 7,884 | 6 | 6 | 6,403 |
| Non-covid | No | Non-covid | 5 | 15,157 | 0 | 5 | 15,157 |
| Asymptomatic | Yes | Asymptomatic | 12 | 33,601 | 1 | 11 | 33,227 |
| Critical | Yes | Critical | 16 | 63,854 | 0 | 16 | 63,854 |
| Mild | Yes | Mild | 19 | 93,835 | 0 | 19 | 93,835 |
| Moderate | Yes | Moderate | 29 | 179,012 | 1 | 28 | 178,688 |
| Severe | Yes | Severe | 7 | 40,235 | 0 | 7 | 40,235 |

**Table 3:**
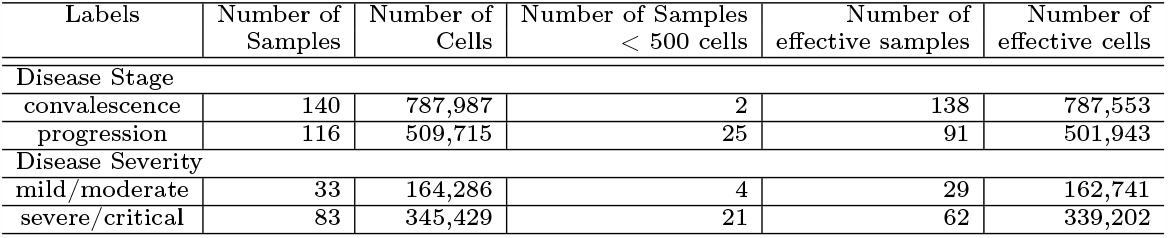
Overview of SC4 scRNA-seq dataset.

| Labels | Number of Samples | Number of Cells | Number of Samples < 500 cells | Number of effective samples | Number of effective cells |
| --- | --- | --- | --- | --- | --- |
| Disease Stage |  |  |  |  |  |
| convalescence | 140 | 787,987 | 2 | 138 | 787,553 |
| progression | 116 | 509,715 | 25 | 91 | 501,943 |
| Disease Severity |  |  |  |  |  |
| mild/moderate | 33 | 164,286 | 4 | 29 | 162,741 |
| severe/critical | 83 | 345,429 | 21 | 62 | 339,202 |

### 2 Evaluations of Phenotype Prediction Accuracy

We provide the precision and recall for different methods on 4 tasks as the complementary information to AUC values described in the main text. Some alternative methods provide better measurements than ScRAT in some specific tasks with some trade-off. For example, Linear provides higher precision in Combat but low recall, and Feedforward (single) provides better recall in most tasks but low precision. These results support that ScRAT provides the best accuracy of phenotype prediction by compromising between precision and recall.

**Figure 1:**
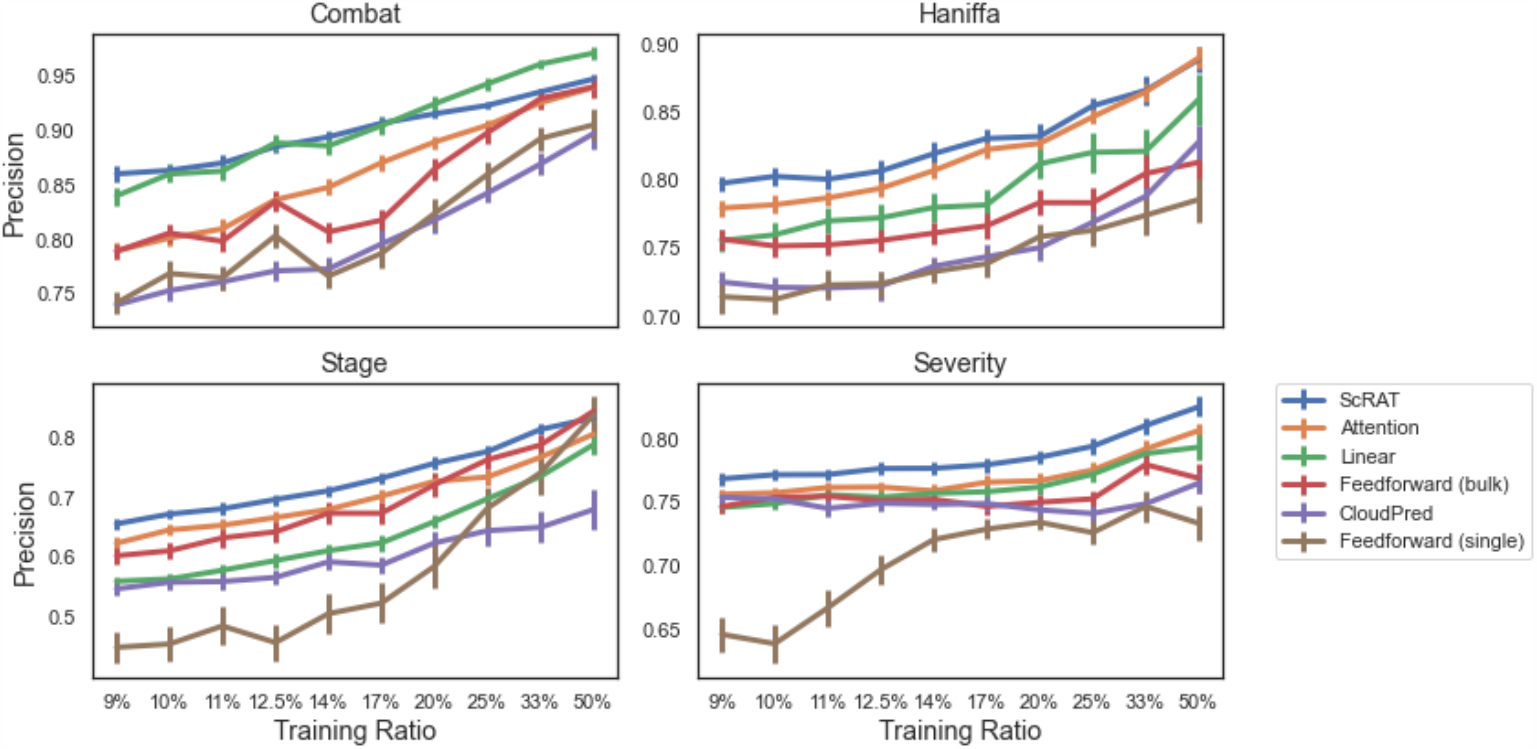
Comparison of different methods on four different tasks based on precision. For each task, we report the prediction accuracy of all methods using precision for 10 different training ratios. ScRAT outperforms other methods in all settings with one exception on the Combat dataset.

### 3 Distribution of High-Attention Cells in the Haniffa Dataset

In addition to the *High-attention Occurrence Value* (HOV) discussed in the main text, we also provide the number of high-attention cells for each cell type of the Haniffa dataset in Table 8. The main differences between this table and the R-score in Table 2 of the main text is that, R-score also considered the occurrence for each cell. Some critical cell types, such as DC3 or CD83_CD14_mono, getting higher rank based on the R-score compared to the one using the ratio of high-attention cells. This indicates that high-attention cells in these cell types are getting high-attention values more often than average. Comprehensive analysis of high-attention cells from these cell types will be the most important goal in the next stage of development of ScRAT.

**Figure 2:**
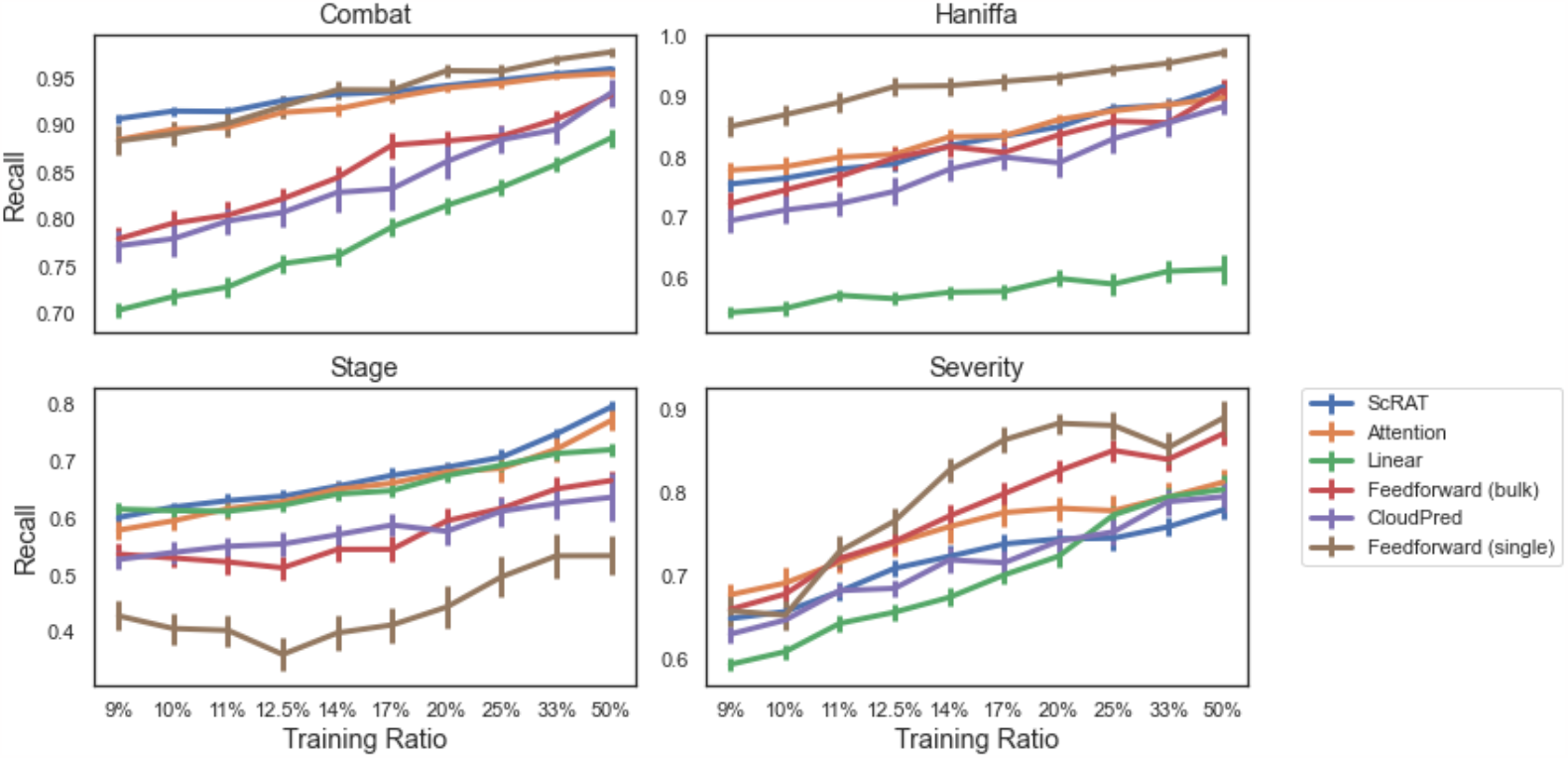
Comparison of different methods on four different tasks based on recall. For each task, we report the prediction accuracy of all methods using recall for 10 different training ratios. Feedforward (single) provides the best recall rates but low precision. This suggests the necessity of simultaneously considering information from all cells to better predict the phenotype of a sample.

**Table 4:** Comparison for different methods on the Haniffa dataset. Best results of AUC, precison, and recall for each training ratio are marked with bold typeface. ScRAT has the highest AUCs in all training ratios while Feedforward (single) provides higher recall with the cost of low precision.

| Haniffa Dataset |  |  |  |  |  |  |  |  |  |  |
| --- | --- | --- | --- | --- | --- | --- | --- | --- | --- | --- |
|  | Training Ratios |  |  |  |  |  |  |  |  |  |
|  | 9% | 10% | 11% | 12.5% | 14% | 17% | 20% | 25% | 33% | 50% |
| Number of Training Samples | 9 | 11 | 12 | 13 | 15 | 18 | 21 | 26 | 35 | 53 |
| AUC |  |  |  |  |  |  |  |  |  |  |
| ScRAT | **0.7532** | **0.7596** | **0.7699** | **0.7806** | **0.8092** | **0.8314** | **0.8456** | **0.8706** | **0.8937** | **0.9207** |
| Attention | 0.7168 | 0.7208 | 0.7368 | 0.7444 | 0.7735 | 0.8009 | 0.8224 | 0.8488 | 0.8816 | 0.9168 |
| Linear | 0.6052 | 0.6119 | 0.6323 | 0.6372 | 0.6543 | 0.6519 | 0.7077 | 0.7126 | 0.7239 | 0.7788 |
| Feedforward (bulk) | 0.6545 | 0.6526 | 0.6597 | 0.6784 | 0.6964 | 0.7031 | 0.7512 | 0.7554 | 0.7971 | 0.8436 |
| CloudPred | 0.6071 | 0.6009 | 0.6175 | 0.6227 | 0.6476 | 0.6681 | 0.6797 | 0.7169 | 0.7675 | 0.8418 |
| Feedforward (single) | 0.6851 | 0.6915 | 0.7023 | 0.7145 | 0.7335 | 0.7414 | 0.7812 | 0.7977 | 0.8290 | 0.8839 |
| Precision |  |  |  |  |  |  |  |  |  |  |
| ScRAT | **0.7980** | **0.8031** | **0.8010** | **0.8070** | **0.8197** | **0.8309** | **0.8324** | **0.8548** | **0.8662** | 0.8888 |
| Attention | 0.7799 | 0.7823 | 0.7873 | 0.7943 | 0.8073 | 0.8228 | 0.8272 | 0.8468 | 0.8650 | **0.8900** |
| Linear | 0.7559 | 0.7602 | 0.7704 | 0.7726 | 0.7803 | 0.7821 | 0.8123 | 0.8207 | 0.8215 | 0.8596 |
| Feedforward (bulk) | 0.7571 | 0.7520 | 0.7528 | 0.7561 | 0.7615 | 0.7668 | 0.7839 | 0.7838 | 0.8052 | 0.8134 |
| CloudPred | 0.7255 | 0.7216 | 0.7213 | 0.7227 | 0.7371 | 0.7441 | 0.7508 | 0.7694 | 0.7884 | 0.8283 |
| Feedforward (single) | 0.7148 | 0.7128 | 0.7234 | 0.7243 | 0.7333 | 0.7391 | 0.7589 | 0.7635 | 0.7745 | 0.7861 |
| Recall |  |  |  |  |  |  |  |  |  |  |
| ScRAT | 0.7566 | 0.7662 | 0.7816 | 0.7899 | 0.8203 | 0.8363 | 0.8512 | 0.8822 | 0.8871 | 0.9178 |
| Attention | 0.7795 | 0.7853 | 0.8009 | 0.8059 | 0.8347 | 0.8364 | 0.8632 | 0.8779 | 0.8877 | 0.8996 |
| Linear | 0.5437 | 0.5506 | 0.5722 | 0.5667 | 0.5771 | 0.5787 | 0.6002 | 0.5909 | 0.6120 | 0.6157 |
| Feedforward (bulk) | 0.7241 | 0.7466 | 0.7694 | 0.8002 | 0.8186 | 0.8088 | 0.8379 | 0.8607 | 0.8580 | 0.9116 |
| CloudPred | 0.6960 | 0.7135 | 0.7242 | 0.7444 | 0.7808 | 0.8012 | 0.7919 | 0.8316 | 0.8579 | 0.8849 |
| Feedforward (single) | **0.8520** | **0.8714** | **0.8921** | **0.9182** | **0.9196** | **0.9266** | **0.9334** | **0.9461** | **0.9568** | **0.9744** |

**Table 5:**
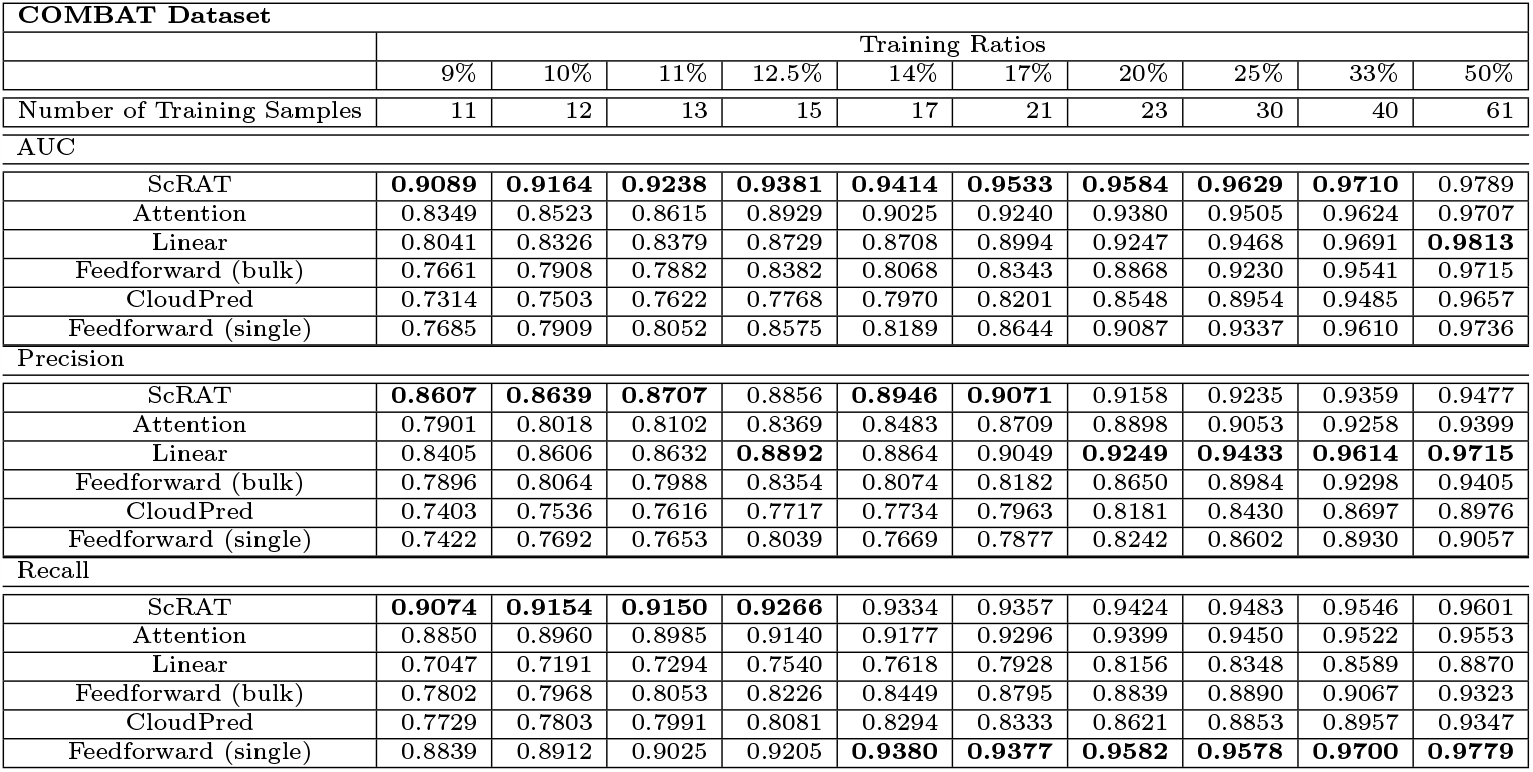
Comparison for different methods on the COMBAT dataset. Best results of AUC, precison, and recall for each training ratio are marked with bold typeface. ScRAT has the highest AUCs in most training ratios while Linear and Feedforward (single) provides higher precision or recall with the cost of low recall/precision.

| COMBAT Dataset |  |  |  |  |  |  |  |  |  |  |
| --- | --- | --- | --- | --- | --- | --- | --- | --- | --- | --- |
|  | Training Ratios |  |  |  |  |  |  |  |  |  |
|  | 9% | 10% | 11% | 12.5% | 14% | 17% | 20% | 25% | 33% | 50% |
| Number of Training Samples | 11 | 12 | 13 | 15 | 17 | 21 | 23 | 30 | 40 | 61 |
| AUC |  |  |  |  |  |  |  |  |  |  |
| ScRAT | **0.9089** | **0.9164** | **0.9238** | **0.9381** | **0.9414** | **0.9533** | **0.9584** | **0.9629** | **0.9710** | 0.9789 |
| Attention | 0.8349 | 0.8523 | 0.8615 | 0.8929 | 0.9025 | 0.9240 | 0.9380 | 0.9505 | 0.9624 | 0.9707 |
| Linear | 0.8041 | 0.8326 | 0.8379 | 0.8729 | 0.8708 | 0.8994 | 0.9247 | 0.9468 | 0.9691 | **0.9813** |
| Feedforward (bulk) | 0.7661 | 0.7908 | 0.7882 | 0.8382 | 0.8068 | 0.8343 | 0.8868 | 0.9230 | 0.9541 | 0.9715 |
| CloudPred | 0.7314 | 0.7503 | 0.7622 | 0.7768 | 0.7970 | 0.8201 | 0.8548 | 0.8954 | 0.9485 | 0.9657 |
| Feedforward (single) | 0.7685 | 0.7909 | 0.8052 | 0.8575 | 0.8189 | 0.8644 | 0.9087 | 0.9337 | 0.9610 | 0.9736 |
| Precision |  |  |  |  |  |  |  |  |  |  |
| ScRAT | **0.8607** | **0.8639** | **0.8707** | 0.8856 | **0.8946** | **0.9071** | 0.9158 | 0.9235 | 0.9359 | 0.9477 |
| Attention | 0.7901 | 0.8018 | 0.8102 | 0.8369 | 0.8483 | 0.8709 | 0.8898 | 0.9053 | 0.9258 | 0.9399 |
| Linear | 0.8405 | 0.8606 | 0.8632 | **0.8892** | 0.8864 | 0.9049 | **0.9249** | **0.9433** | **0.9614** | **0.9715** |
| Feedforward (bulk) | 0.7896 | 0.8064 | 0.7988 | 0.8354 | 0.8074 | 0.8182 | 0.8650 | 0.8984 | 0.9298 | 0.9405 |
| CloudPred | 0.7403 | 0.7536 | 0.7616 | 0.7717 | 0.7734 | 0.7963 | 0.8181 | 0.8430 | 0.8697 | 0.8976 |
| Feedforward (single) | 0.7422 | 0.7692 | 0.7653 | 0.8039 | 0.7669 | 0.7877 | 0.8242 | 0.8602 | 0.8930 | 0.9057 |
| Recall |  |  |  |  |  |  |  |  |  |  |
| ScRAT | **0.9074** | **0.9154** | **0.9150** | **0.9266** | 0.9334 | 0.9357 | 0.9424 | 0.9483 | 0.9546 | 0.9601 |
| Attention | 0.8850 | 0.8960 | 0.8985 | 0.9140 | 0.9177 | 0.9296 | 0.9399 | 0.9450 | 0.9522 | 0.9553 |
| Linear | 0.7047 | 0.7191 | 0.7294 | 0.7540 | 0.7618 | 0.7928 | 0.8156 | 0.8348 | 0.8589 | 0.8870 |
| Feedforward (bulk) | 0.7802 | 0.7968 | 0.8053 | 0.8226 | 0.8449 | 0.8795 | 0.8839 | 0.8890 | 0.9067 | 0.9323 |
| CloudPred | 0.7729 | 0.7803 | 0.7991 | 0.8081 | 0.8294 | 0.8333 | 0.8621 | 0.8853 | 0.8957 | 0.9347 |
| Feedforward (single) | 0.8839 | 0.8912 | 0.9025 | 0.9205 | **0.9380** | **0.9377** | **0.9582** | **0.9578** | **0.9700** | **0.9779** |

**Table 6:**
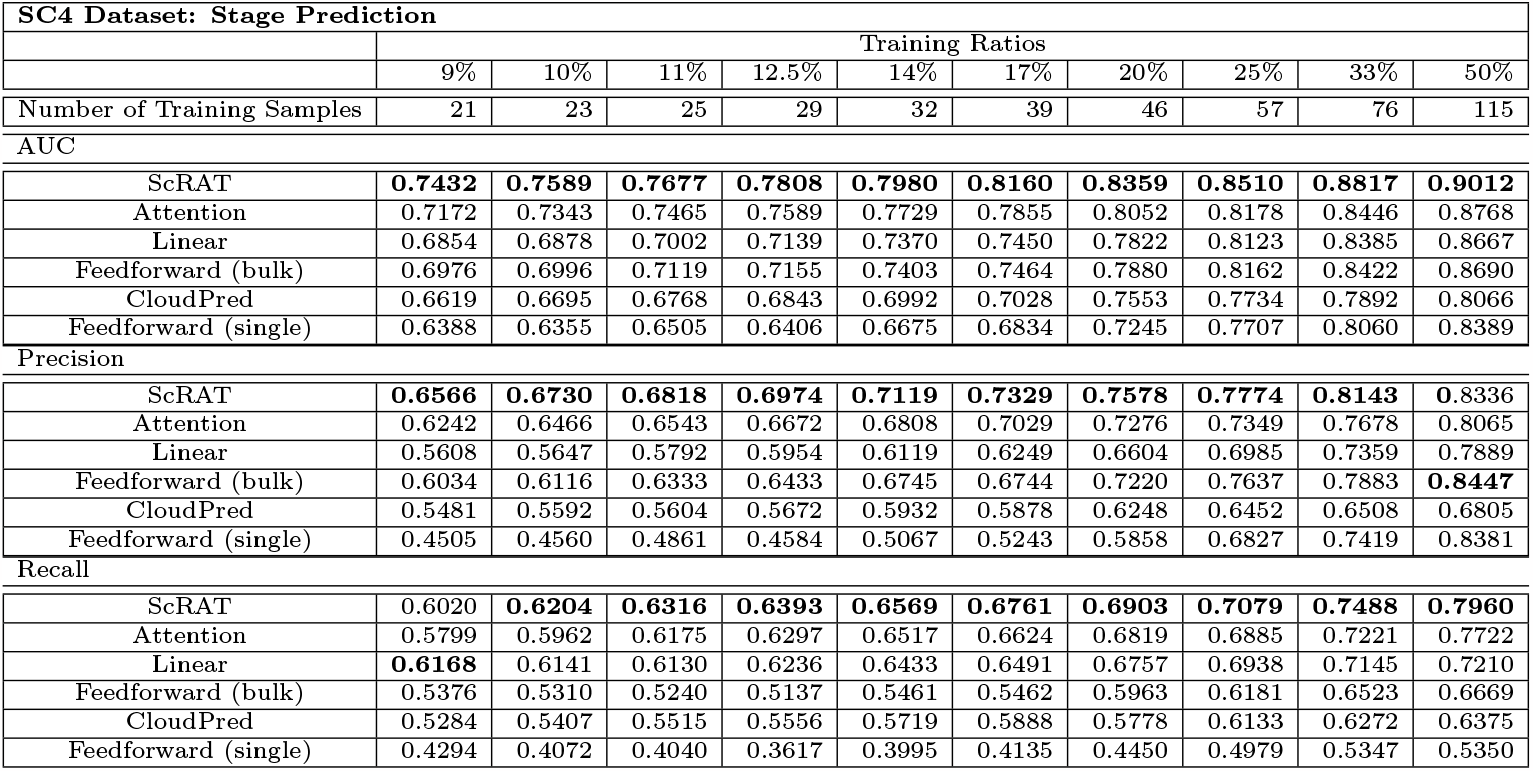
Comparison for different methods on the Stage prediction task for SC4 dataset. Best results of AUC, precison, and recall for each training ratio are marked with bold typeface. ScRAT has the highest AUCs in all training ratios and also achieve best precision and recall for most training ratios.

**Table 7:** Comparison for different methods on the Severity prediction task for SC4 dataset. Best results of AUC, precison, and recall for each training ratio are marked with bold typeface. ScRAT has the highest AUCs and precision in all training ratios while Attenion and Feedforward (single) provides higher recall with the cost of low precision.

| SC4 Dataset: Severity Prediction |  |  |  |  |  |  |  |  |  |  |
| --- | --- | --- | --- | --- | --- | --- | --- | --- | --- | --- |
|  | Training Ratios |  |  |  |  |  |  |  |  |  |
|  | 9% | 10% | 11% | 12.5% | 14% | 17% | 20% | 25% | 33% | 50% |
| Number of Training Samples | 8 | 9 | 10 | 11 | 13 | 15 | 18 | 23 | 30 | 46 |
| AUC |  |  |  |  |  |  |  |  |  |  |
| ScRAT | **0.6091** | **0.6124** | **0.6185** | **0.6328** | **0.6352** | **0.6450** | **0.6644** | **0.6800** | **0.7038** | **0.7410** |
| Attention | 0.5986 | 0.6064 | 0.6137 | 0.6242 | 0.6174 | 0.6374 | 0.6485 | 0.6709 | 0.7028 | 0.7197 |
| Linear | 0.5672 | 0.5767 | 0.5957 | 0.6012 | 0.6108 | 0.6214 | 0.6379 | 0.6668 | 0.6986 | 0.7007 |
| Feedforward (bulk) | 0.5901 | 0.6016 | 0.6102 | 0.6170 | 0.6251 | 0.6315 | 0.6425 | 0.6646 | 0.6993 | 0.6898 |
| CloudPred | 0.5819 | 0.6078 | 0.6069 | 0.6001 | 0.6140 | 0.6176 | 0.6114 | 0.6337 | 0.6446 | 0.6541 |
| Feedforward (single) | 0.5778 | 0.5869 | 0.5930 | 0.5897 | 0.5941 | 0.5911 | 0.5983 | 0.6103 | 0.6620 | 0.6422 |
| Precision |  |  |  |  |  |  |  |  |  |  |
| ScRAT | **0.7689** | **0.7721** | **0.7722** | **0.7769** | **0.7771** | **0.7799** | **0.7857** | **0.7944** | **0.8106** | **0.8252** |
| Attention | 0.7568 | 0.7580 | 0.7621 | 0.7624 | 0.7595 | 0.7662 | 0.7674 | 0.7757 | 0.7921 | 0.8066 |
| Linear | 0.7465 | 0.7494 | 0.7564 | 0.7545 | 0.7575 | 0.7588 | 0.7625 | 0.7724 | 0.7886 | 0.7936 |
| Feedforward (bulk) | 0.7472 | 0.7543 | 0.7553 | 0.7518 | 0.7528 | 0.7473 | 0.7505 | 0.7532 | 0.7801 | 0.7693 |
| CloudPred | 0.7544 | 0.7532 | 0.7458 | 0.7497 | 0.7489 | 0.7495 | 0.7446 | 0.7418 | 0.7493 | 0.7654 |
| Feedforward (single) | 0.6472 | 0.6398 | 0.6679 | 0.6977 | 0.7213 | 0.7295 | 0.7349 | 0.7268 | 0.7472 | 0.7340 |
| Recall |  |  |  |  |  |  |  |  |  |  |
| ScRAT | 0.6499 | 0.6582 | 0.6825 | 0.7099 | 0.7242 | 0.7390 | 0.7450 | 0.7460 | 0.7595 | 0.7798 |
| Attention | **0.6784** | **0.6923** | 0.7175 | 0.7401 | 0.7597 | 0.7765 | 0.7817 | 0.7788 | 0.7950 | 0.8127 |
| Linear | 0.5953 | 0.6104 | 0.6443 | 0.6576 | 0.6753 | 0.7020 | 0.7244 | 0.7736 | 0.7953 | 0.8040 |
| Feedforward (bulk) | 0.6610 | 0.6792 | 0.7218 | 0.7421 | 0.7723 | 0.7990 | 0.8262 | 0.8507 | 0.8400 | 0.8705 |
| CloudPred | 0.6314 | 0.6481 | 0.6833 | 0.6860 | 0.7200 | 0.7166 | 0.7424 | 0.7534 | 0.7895 | 0.7953 |
| Feedforward (single) | 0.6591 | 0.6540 | **0.7306** | **0.7664** | **0.8270** | **0.8632** | **0.8830** | **0.8805** | **0.8540** | **0.8892** |

**Table 8:**
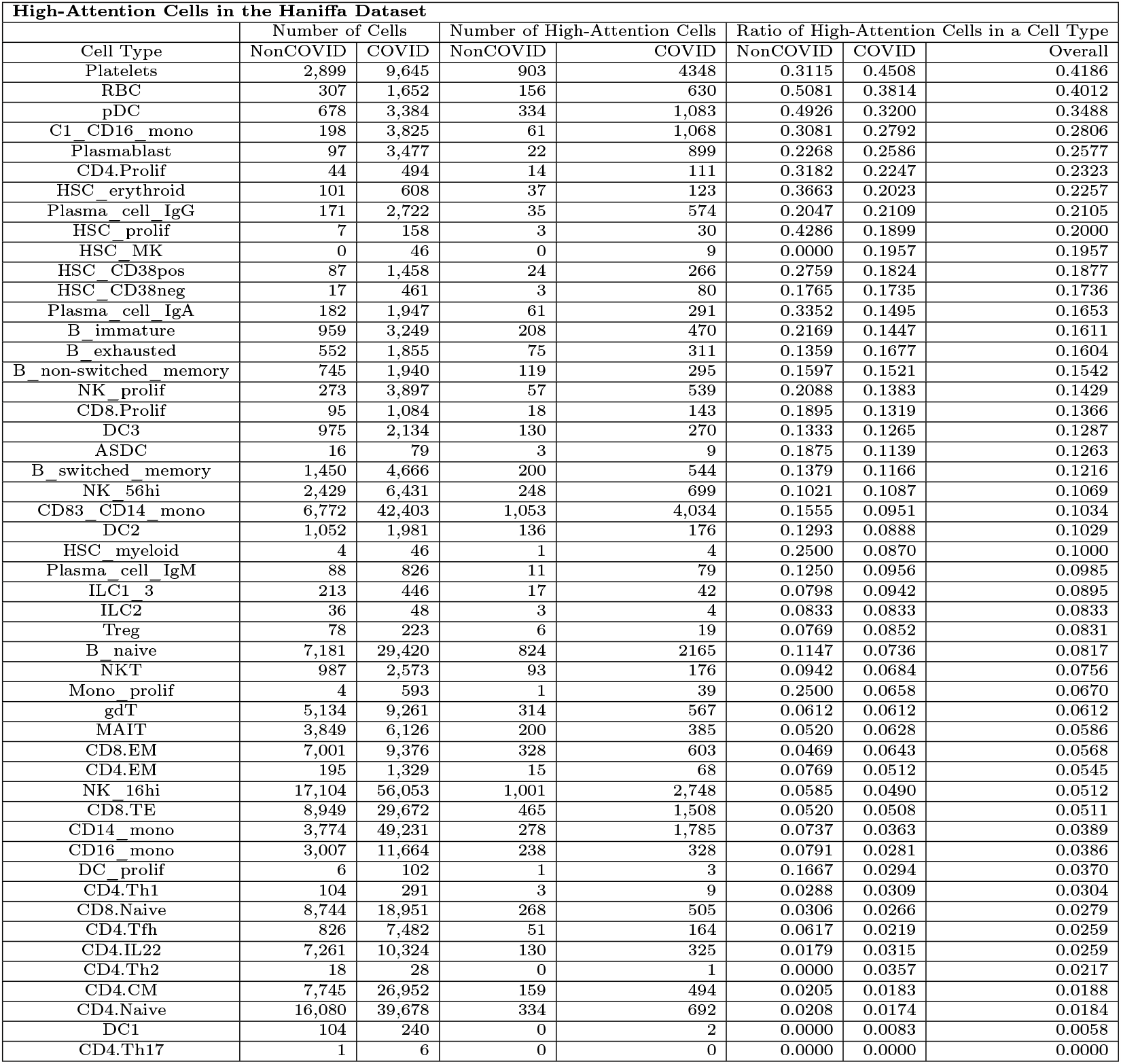
Number of high-attention cells for each cell type in the Haniffa dataset. Compared to the HOVs that weight each cell using the high-attention *Occurrence*, here we only count the number of high-attention cells for each cell type. In other words, we force the HOVs = 1 for all high-attention cells to understand their distribution. Note that the *NonCOVID, COVID*, and *Overall* values in the “Ratio of High-Attention Cells in a Cell Type” section correspond to the ratio of high-attention cells in a cell type for NonCOVID patients, COVID patients, and all patients respectively. The differences between the rankings based on these ratios and the R-score (in Table 2 of the main text) suggest that some specific cell types include cells getting high-attention values more often than average. More analysis of these cells has the potential to improve our understanding of attention mechanisms in the domain of single-cell biology.

| High-Attention Cells in the Haniffa Dataset |  |  |  |  |  |  |  |
| --- | --- | --- | --- | --- | --- | --- | --- |
| Cell Type | Number of Cells |  | Number of High-Attention Cells |  | Ratio of High-Attention Cells in a Cell Type |  |  |
|  | NonCOVID | COVID | NonCOVID | COVID | NonCOVID | COVID | Overall |
| Platelets | 2,899 | 9,645 | 903 | 4348 | 0.3115 | 0.4508 | 0.4186 |
| RBC | 307 | 1,652 | 156 | 630 | 0.5081 | 0.3814 | 0.4012 |
| pDC | 678 | 3,384 | 334 | 1,083 | 0.4926 | 0.3200 | 0.3488 |
| C1_CD16_mono | 198 | 3,825 | 61 | 1,068 | 0.3081 | 0.2792 | 0.2806 |
| Plasmablast | 97 | 3,477 | 22 | 899 | 0.2268 | 0.2586 | 0.2577 |
| CD4.Prolif | 44 | 494 | 14 | 111 | 0.3182 | 0.2247 | 0.2323 |
| HSC_erythroid | 101 | 608 | 37 | 123 | 0.3663 | 0.2023 | 0.2257 |
| Plasma_cell_IgG | 171 | 2,722 | 35 | 574 | 0.2047 | 0.2109 | 0.2105 |
| HSC_prolif | 7 | 158 | 3 | 30 | 0.4286 | 0.1899 | 0.2000 |
| HSC_MK | 0 | 46 | 0 | 9 | 0.0000 | 0.1957 | 0.1957 |
| HSC_CD38pos | 87 | 1,458 | 24 | 266 | 0.2759 | 0.1824 | 0.1877 |
| HSC_CD38neg | 17 | 461 | 3 | 80 | 0.1765 | 0.1735 | 0.1736 |
| Plasma_cell_IgA | 182 | 1,947 | 61 | 291 | 0.3352 | 0.1495 | 0.1653 |
| B_immature | 959 | 3,249 | 208 | 470 | 0.2169 | 0.1447 | 0.1611 |
| B_exhausted | 552 | 1,855 | 75 | 311 | 0.1359 | 0.1677 | 0.1604 |
| B_non-switched_memory | 745 | 1,940 | 119 | 295 | 0.1597 | 0.1521 | 0.1542 |
| NK_prolif | 273 | 3,897 | 57 | 539 | 0.2088 | 0.1383 | 0.1429 |
| CD8.Prolif | 95 | 1,084 | 18 | 143 | 0.1895 | 0.1319 | 0.1366 |
| DC3 | 975 | 2,134 | 130 | 270 | 0.1333 | 0.1265 | 0.1287 |
| ASDC | 16 | 79 | 3 | 9 | 0.1875 | 0.1139 | 0.1263 |
| B_switched_memory | 1,450 | 4,666 | 200 | 544 | 0.1379 | 0.1166 | 0.1216 |
| NK_56hi | 2,429 | 6,431 | 248 | 699 | 0.1021 | 0.1087 | 0.1069 |
| CD83_CD14_mono | 6,772 | 42,403 | 1,053 | 4,034 | 0.1555 | 0.0951 | 0.1034 |
| DC2 | 1,052 | 1,981 | 136 | 176 | 0.1293 | 0.0888 | 0.1029 |
| HSC_myeloid | 4 | 46 | 1 | 4 | 0.2500 | 0.0870 | 0.1000 |
| Plasma_cell_IgM | 88 | 826 | 11 | 79 | 0.1250 | 0.0956 | 0.0985 |
| ILC1_3 | 213 | 446 | 17 | 42 | 0.0798 | 0.0942 | 0.0895 |
| ILC2 | 36 | 48 | 3 | 4 | 0.0833 | 0.0833 | 0.0833 |
| Treg | 78 | 223 | 6 | 19 | 0.0769 | 0.0852 | 0.0831 |
| B_naive | 7,181 | 29,420 | 824 | 2165 | 0.1147 | 0.0736 | 0.0817 |
| NKT | 987 | 2,573 | 93 | 176 | 0.0942 | 0.0684 | 0.0756 |
| Mono_prolif | 4 | 593 | 1 | 39 | 0.2500 | 0.0658 | 0.0670 |
| gdT | 5,134 | 9,261 | 314 | 567 | 0.0612 | 0.0612 | 0.0612 |
| MAIT | 3,849 | 6,126 | 200 | 385 | 0.0520 | 0.0628 | 0.0586 |
| CD8.EM | 7,001 | 9,376 | 328 | 603 | 0.0469 | 0.0643 | 0.0568 |
| CD4.EM | 195 | 1,329 | 15 | 68 | 0.0769 | 0.0512 | 0.0545 |
| NK_16hi | 17,104 | 56,053 | 1,001 | 2,748 | 0.0585 | 0.0490 | 0.0512 |
| CD8.TE | 8,949 | 29,672 | 465 | 1,508 | 0.0520 | 0.0508 | 0.0511 |
| CD14_mono | 3,774 | 49,231 | 278 | 1,785 | 0.0737 | 0.0363 | 0.0389 |
| CD16_mono | 3,007 | 11,664 | 238 | 328 | 0.0791 | 0.0281 | 0.0386 |
| DC_prolif | 6 | 102 | 1 | 3 | 0.1667 | 0.0294 | 0.0370 |
| CD4.Th1 | 104 | 291 | 3 | 9 | 0.0288 | 0.0309 | 0.0304 |
| CD8.Naive | 8,744 | 18,951 | 268 | 505 | 0.0306 | 0.0266 | 0.0279 |
| CD4.Tfh | 826 | 7,482 | 51 | 164 | 0.0617 | 0.0219 | 0.0259 |
| CD4.IL22 | 7,261 | 10,324 | 130 | 325 | 0.0179 | 0.0315 | 0.0259 |
| CD4.Th2 | 18 | 28 | 0 | 1 | 0.0000 | 0.0357 | 0.0217 |
| CD4.CM | 7,745 | 26,952 | 159 | 494 | 0.0205 | 0.0183 | 0.0188 |
| CD4.Naive | 16,080 | 39,678 | 334 | 692 | 0.0208 | 0.0174 | 0.0184 |
| DC1 | 104 | 240 | 0 | 2 | 0.0000 | 0.0083 | 0.0058 |
| CD4.Th17 | 1 | 6 | 0 | 0 | 0.0000 | 0.0000 | 0.0000 |

